# AlphaPart - R implementation of the method for partitioning genetic trends

**DOI:** 10.1101/2020.04.24.059071

**Authors:** Jana Obšteter, Justin Holl, John M. Hickey, Gregor Gorjanc

## Abstract

**Background:** In this paper we present the AlphaPart R package, an open-source software that implements a method for partitioning breeding values and genetic trends to identify sources of genetic gain. Breeding programmes improve populations for a set of traits, which can be measured with a genetic trend calculated from averaged year of birth estimated breeding values of selection candidates. While sources of the overall genetic gain are generally known, their realised contributions are hard to quantify in complex breeding programmes. The aim of this paper is to present the AlphaPart R package and demonstrate it with a simulated pig breeding example.

**Results:** The package includes the main partitioning function AlphaPart, that partitions the breeding values and genetic trends by analyst defined paths, and a set of functions for handling data and results. The package is freely available from CRAN repository at http://CRAN.R-project.org/package=AlphaPart. We demonstrate the use of the package by examining the genetic gain in a pig breeding example, in which the multiplier achieved higher breeding values than the nucleus for traits measured and selected in the multiplier. The partitioning analysis revealed that these higher values depended on the accuracy and intensity of selection in the multiplier and the extent of gene flow from the nucleus. For traits measured only in the nucleus, the multiplier achieved comparable or smaller genetic gain than the nucleus depending on the amount of gene flow.

**Conclusions:** AlphaPart implements a method for partitioning breeding values and genetic trends and provides a useful tool for quantifying the sources of genetic gain in breeding programmes. The use of AlphaPart will help breeders to better understand or improve their breeding programmes.

## Background

In this paper we present the AlphaPart R package that implements a method for partitioning breeding values and genetic trends, and demonstrate it with a pig breeding example. Breeding programmes improve populations for a set of traits by selecting and intermating genetically superior individuals. Population improvement can be measured with a genetic trend calculated from averaged year of birth estimated breeding values of selection candidates [1,2].

While sources of the overall genetic gain are generally known, their realised contributions are hard to quantify in complex breeding programmes. García-Cortés *et al*. [3] proposed a method for such analysis. First, the method partitions breeding values into parent average and Mendelian sampling terms [4], and allocates the terms to analyst-defined “paths” (males, females, tested sires, etc.). Next, it summarizes path specific terms to quantify path contributions to the overall genetic trend.

The partitioning method has been used in a number of cases. Gorjanc *et al*. [5] and Gorjanc *et al*. [6] estimated contributions of national breeding programmes to Brows-Swiss and Holstein country-specific and global genetic trends. Špehar *et al*. [7] estimated contributions of national selection and importation in Croatian Simmental cattle. Škorput *et al*. [8] estimated the contribution of national selection and importation in two pig breeds in Croatia, and extended the analysis with the quantification of uncertainty [2]. However, these studies used dedicated software implementations of the partitioning method, for which no open-source software exists.

The aim of this paper is to: i) present the AlphaPart R package; and ii) demonstrate it with a simulated pig breeding example that quantifies nucleus-multiplier gene flow and the contribution of nucleus and multiplier selection on genetic gain in the two tiers.

## Implementation

We first demonstrate the AlphaPart package and its functions on an example dataset. Next, we describe the simulation of a pig breeding example to demonstrate the use of AlphaPart.

### AlphaPart

AlphaPart is an R package available from CRAN repository at https://CRAN.R-project.org/package=AlphaPart. It consists of the main function `AlphaPart` for partitioning breeding values and auxiliary functions for manipulating data and summarizing, visualizing, and saving results. The package includes an example dataset `AlphaPart.ped`, which includes a four-generation pedigree and information about the generation, country, gender, and breeding values. Below we describe and demonstrate the functions with the dataset.

We install and load the package with:

~~~
> install.packages(pkg = “AlphaPart”)
> library(package = “AlphaPart”)
~~~

We use the `AlphaPart` function to partition breeding values (`bv1`) in the `AlphaPart.ped` by the country variable into domestic and import contributions:

~~~
> data(AlphaPart.ped)
> part <- AlphaPart(x = AlphaPart.ped,
                        colPath = “country”,
                        colBV = “bv1”)
~~~

The partitioning function `AlphaPart` requires a data frame holding pedigree with animal/sire/dam or animal/sire/maternal-grandsire, a time-ordering variable such as year of birth, partition variable (path), and breeding values. Following the method described in García-Cortés *et al*. [3], we recurse the pedigree from the oldest to the youngest individual, for each individual calculate parent average and Mendelian sampling terms for any number of traits and assign terms to paths. We partition multiple traits by specifying a vector of variables, say `colBV = c(“bv1”, “bv2”)`. The multiple trait option can also serve to partition samples from a posterior distribution to quantify uncertainty [2, 8]. To speed-up calculations we use C++ and trait-vectorised partitioning. The function can also directly partition and summarize path contributions “on-the-fly”, which is a useful computational speed-up for huge pedigrees. The output object of the function is either `AlphaPart` or `summaryAlphaPart` class.

We use the generic `summary.AlphaPar`t function to summarize an `AlphaPart` object by a grouping variable, say generation (`gen`):

~~~
> sumPartByGen <- summary(part, by = “gen”)
> print(sumPartByGen)
~~~

The `summary` function summarizes breeding values and their path partitions by levels of grouping variable. By default, we summarize with a mean, but the user can specify any R function via the `FUN` argument. The `summary` function can also summarize only a subset of the object via the `subset` argument.

We use the generic `plot.summaryAlphaPart` function to plot summarized partitions:

~~~
> plot(sumPartByGen)
~~~

We also provide a number of utility functions that ease partitioning analysis. With the `pedFixBirthYear` function we impute missing or fix erroneous years of birth. With the `pedSetBase` function we set the base population by specifying founders and removing older pedigree records. With the `AlphaPartSubset` function we keep partitions for specified paths in the `AlphaPart` or `summaryAlphaPart` objects. With the `AlphaPartSum` function we sum the partitions of several paths in a `summaryAlphaPart` object. The `AlphaPartSubset` and `AlphaPartSum` functions simplify the presentation of partitioning analysis.

### Pig breeding example

We applied the AlphaPart R package to a simulated pig breeding example to examine the nucleus-multiplier gene flow and the contribution of nucleus and multiplier selection on genetic gain in both tiers. Pig breeders select in the nucleus and multiply this improvement in the multiplier to supply large number of commercial animals. The multiplier generally has lower genetic mean than the nucleus due to time-lag. However, animals with very high breeding values are often observed in the multiplier for some traits and we aimed to use AlphaPart to explain the source of this observation. To this end we have first simulated a stylised pig breeding programme that exposes the drivers of real observations. We have next partitioned the genetic trend of true breeding values by a tier-gender variable to quantify sources of genetic gain in the nucleus and the multiplier.

We used the AlphaSimR package [9] to simulate a pig breeding programme for a single breed with 40 years of selection on two uncorrelated traits. Trait 1 had heritability 0.25 and trait 2 had heritability 0.10. We measured both traits in the nucleus and only trait 1 in the multiplier.

We selected on the index of the two traits with equal emphasis. We split the simulation into initial 20 years of a burn-in and 20 years of evaluation.

In the burn-in we simulated only the nucleus and selected animals based on the index of phenotype values for both traits. We selected 25 males and 500 females each year and randomly crossed them to produce a new generation of 6,000 progeny (12 per cross). At the end of the burn-in we generated 5,000 females to seed the multiplier.

In the evaluation we simulated both the nucleus and the multiplier and selected animals based on the index of estimated breeding values for both traits. In the nucleus, we selected 25 males and 500 females each year and randomly crossed them to produce a new generation of 6,000 progeny (12 per cross). In the multiplier, we selected 750 females each year and randomly crossed them to a set of males to produce a new generation of 9,000 progeny (12 per cross). To quantify the effect of selection in the multiplier on genetic gain we defined the set of males as either 1) the 25 best nucleus males (MaleFlow100 scenario) or 2) the 25 best nucleus males and 100 best multiplier males (MaleFlow20 scenario).

We estimated the breeding values for each trait independently before each nucleus or multiplier selection decision. We ran pedigree-based model implemented in blupf90 [10] and used all available data from evaluation years. The model included the mean as a fixed effect and animal breeding values as a random effect modelled hierarchically with pedigree.

Finally, we partitioned the true breeding values and genetic trends with the AlphaPart as demonstrated above. We used `AlphaPart` function to partition true breeding values from the 20 evaluation years by the tier-gender variable and `summary.AlphaPart` function to summarize the partitions by year to quantify the contribution of each tier-gender level to genetic trend in the nucleus and the multiplier.

We repeated the simulation 10 times. We present standardized true breeding values and genetic trends, as well as their partitions with mean set to zero and genetic standard deviation set to one in the year 20. We chose to present true (instead of estimated) breeding values to assess the true sources of genetic gain. The simulation code for the datasets generated and/or analysed during the current study are available in the GitLab repository, https://git.ecdf.ed.ac.uk/HighlanderLab_public/jobsteter_alphapart.

## Results

The results show partitions of true breeding values and genetic trends obtained with the AlphaPart for the two simulated pig breeding scenarios. Partitioning showed that we can explain the situation with very high breeding values in the multiplier by the extent of nucleus-multiplier gene flow as well as accuracy and intensity of multiplier selection.

### Partitioning the true breeding values and genetic trend of MaleFlow100 scenario

In MaleFlow100 scenario the multiplier achieved a higher final genetic gain than the nucleus for trait 1 due to selection of multiplier females. We show this in Figure 1 that presents the distribution of true breeding values and their partitions by trait and tier for two years of one replicate of MaleFlow100 scenario, and in Figure 2 that presents the genetic trends and their partitions summarised over 10 replicates. The partitioning expectedly showed that in the nucleus the genetic gain stemmed from selection of nucleus males and nucleus females. However, the contribution of male and female selection changed over the years. While in year 23 the contributions of male and female selection were more comparable, by year 40 male selection contributed more. The mean final genetic gain in the nucleus for trait 1 was 9.75 and 8.34 for trait 2, with male selection contributing 5.65 for trait 1 and 4.92 for trait 2, and female selection contributing 4.10 for trait 1 and 3.42 for trait 2.

**Figure 1.**
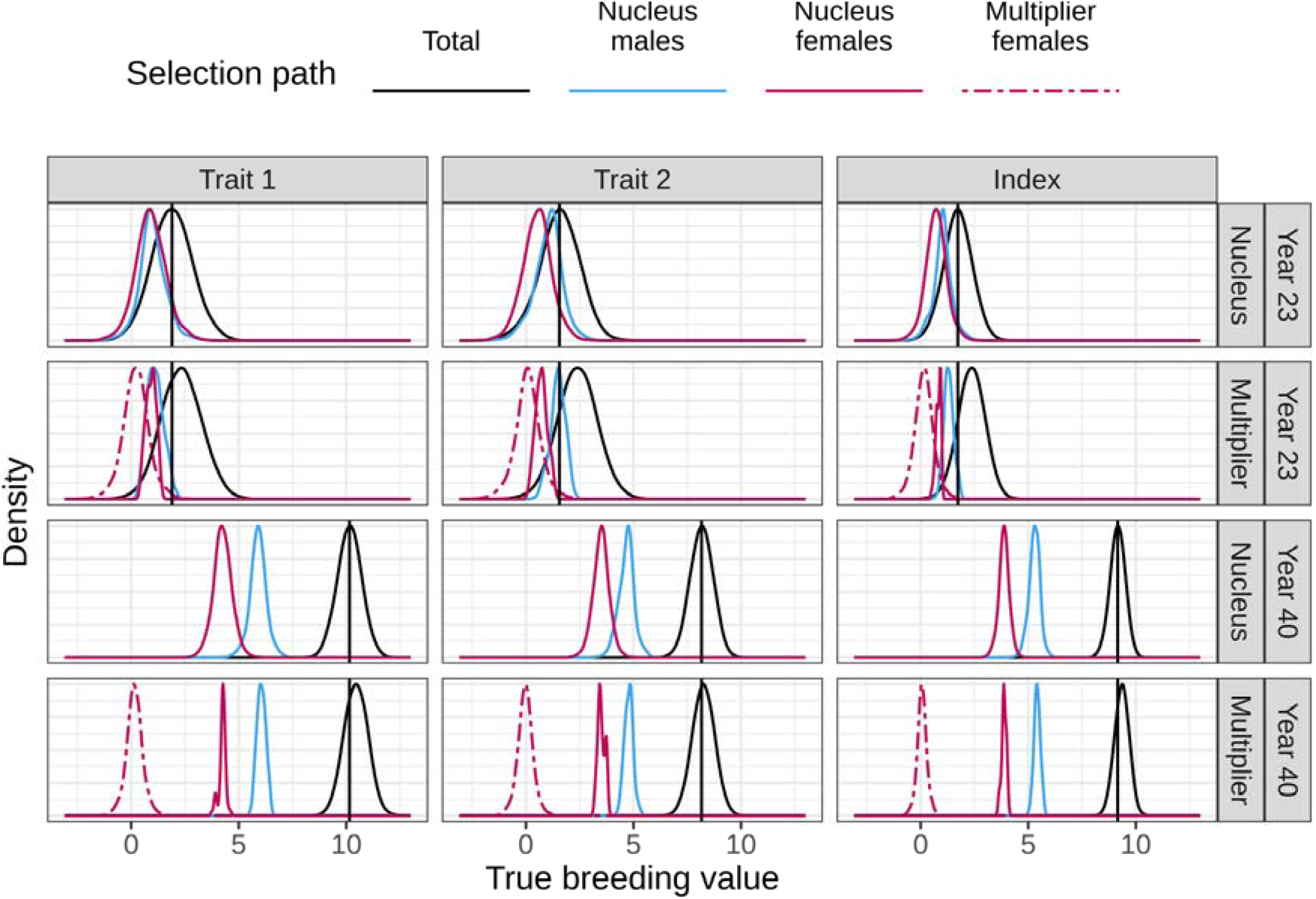
Distribution of true breeding values and their partitions by trait, year, and tier in MaleFlow100 scenario. We show scaled densities of partitions in years 23 and 40 of one simulation replicate. MalerFlow100 uses only nucleus males in the multiplier. Trait 1 is measured in the nucleus and the multiplier, while trait 2 is measured only in the nucleus. Black vertical lines represent the nucleus mean breeding value for a trait in a year.

**Figure 2:**
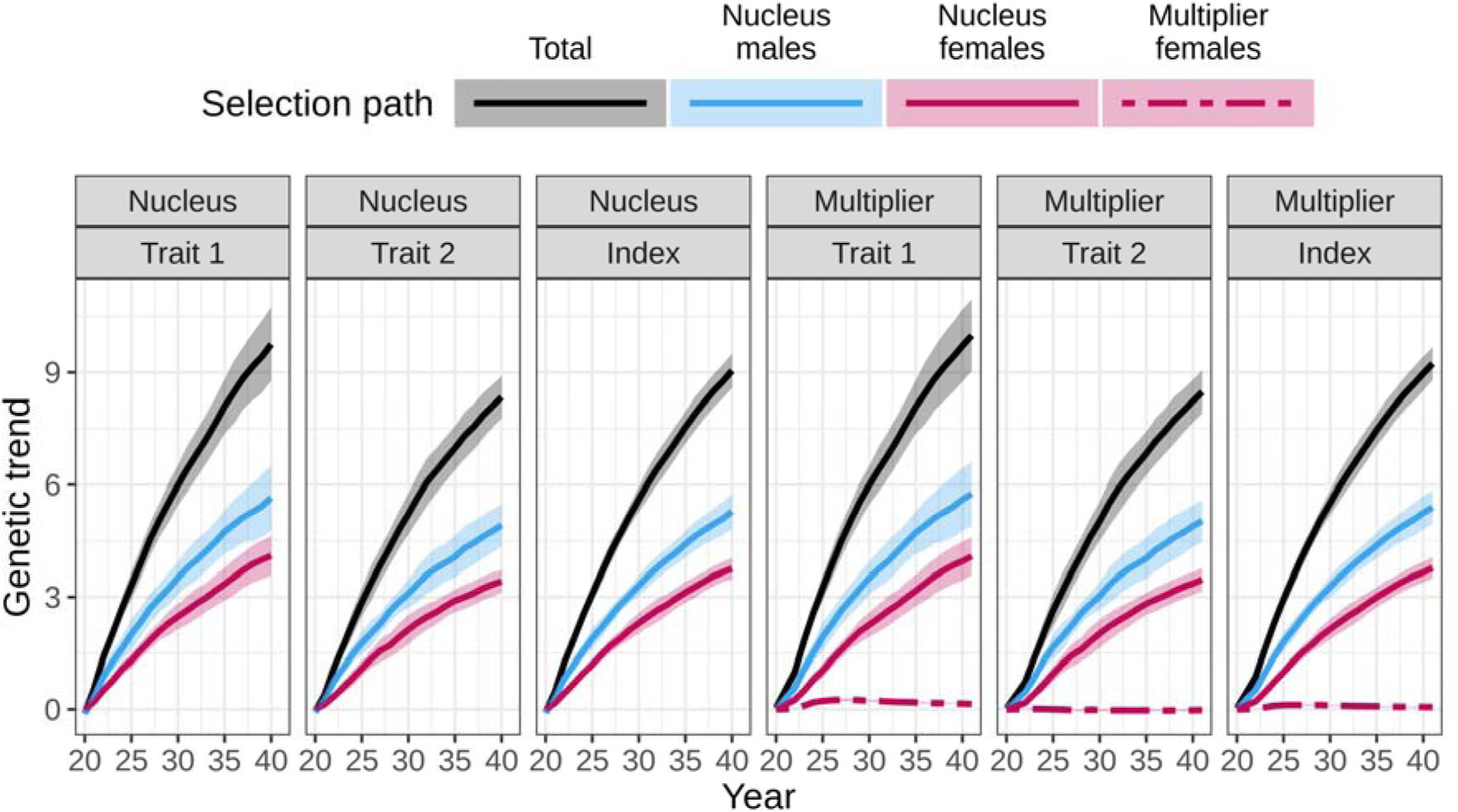
Partitioning of genetic trend by tier-gender in MaleFlow100 scenario. The scenario uses nucleus males in the multiplier. Trait 1 is measured in the nucleus and the multiplier, while trait 2 is measured only in the nucleus.

In the multiplier, the genetic gain was higher than in the nucleus. In year 23 the multiplier had higher genetic gain than the nucleus for both traits, while in year 40 it had higher genetic gain only for trait 1. The higher genetic gain was partly due to larger contribution of nucleus males in multiplier than in nucleus (via gene flow) and partly due to non-zero contribution of multiplier female selection. The mean final genetic gain in the multiplier for trait 1 was 10.00 with nucleus males contributing 5.75, nucleus females 4.09, and multiplier females 0.14. The mean final genetic gain and path partitions for trait 2 in the multiplier were comparable to the nucleus.

### Partitioning the true breeding values and genetic trend of MaleFlow20 scenario

In the MaleFlow20 scenario selection of multiplier males further increased the final genetic gain for trait 1 in the multiplier compared to the nucleus, but decreased the final genetic gain for trait 2. We show this in Figure 3 that presents the distribution of true breeding values and their partitions by trait and tier for two year of one replicate of MaleFlow20 scenario, and Figure 4 that presents the genetic trends and their partitions summarised over 10 replicates. As in MaleFlow100 scenario, the nucleus genetic gain stemmed from selection of nucleus males and females. Progressing from year 23 to year 40, the contribution of nucleus males increased compared to nucleus females, but only for trait 1. The mean final genetic gain for trait 1 was 10.09 and 8.39 for trait 2, with nucleus males contributing 5.69 for trait 1 and 5.17 for trait 2, and nucleus females contributing 4.40 for trait 1 and 3.22 for trait 2.

**Figure 3.**
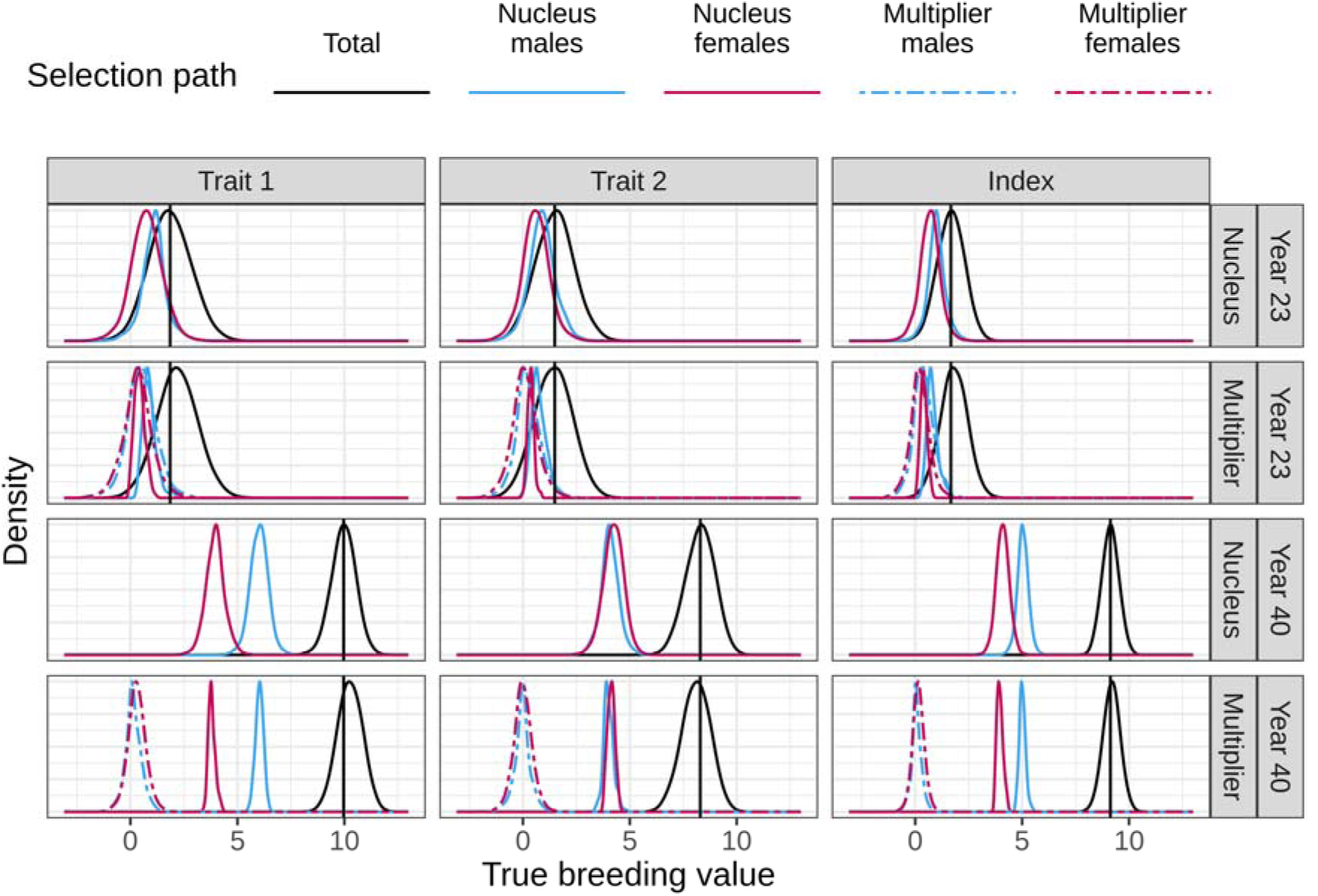
Distribution of true breeding values and their partitions by trait, year, and tier in MaleFlow20 scenario. We show scaled densities of partitions in years 23 and 40 of one simulation replicate. MalerFlow20 uses nucleus and multiplier males in the multiplier. Trait 1 is measured in the nucleus and the multiplier, while trait 2 is measured only in the nucleus. Black vertical lines represent the nucleus mean breeding value for a trait in a year.

**Figure 4.**
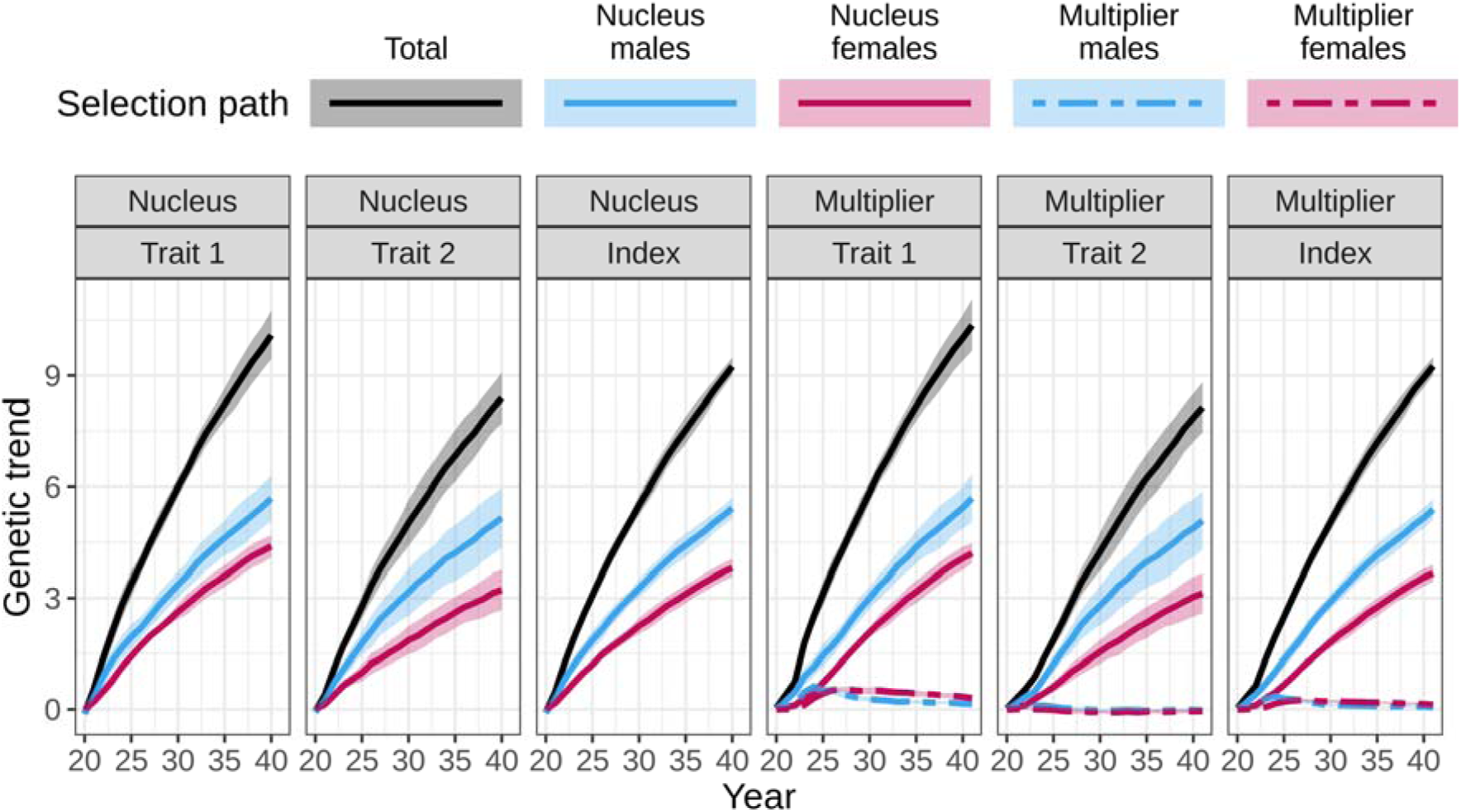
Partitioning of the genetic trend by tier-gender in MaleFlow20 scenario. The scenarios uses nucleus and multiplier males in the multiplier. Trait 1 is measured in the nucleus and the multiplier, while trait 2 is measured only in the nucleus.

In the multiplier the genetic gain was again higher than in the nucleus, but only for trait 1. This higher genetic gain was consistent throughout the generations and was a result of selection of multiplier females and multiplier males, and reduced contribution of gene flow from the nucleus females (via reduced use of nucleus males). For trait 2, the multiplier performed progressively worse relative to the nucleus over the years. In year 40, the genetic gain in the multiplier was lower than in the nucleus due to a small average negative contribution of multiplier females and multiplier males and reduced contribution of gene flow from the nucleus females and nucleus males. The mean final genetic gain in the multiplier was 10.36 for trait 1 and 8.14 for trait 2, with nucleus males contributing 5.70 for trait 1 and 5.09 for trait 2, nucleus females contributing 4.21 for trait 1 and 3.13 for trait 2, multiplier males contributing 0.15 for trait 1 and -0.03 for trait 2, and multiplier females contributing 0.30 for trait 1 and -0.05 for trait 2.

## Discussion

In this paper we present AlphaPart, freely available R package that implements the method for partitioning breeding values and genetic trends. We demonstrate the package on a simulated pig breeding example with a higher genetic trend for some traits in the multiplier compared to the nucleus. Following this, we organized the discussion into two parts: i) advantages and disadvantages of the AlphaPart R package; ii) partitioning results of the pig breeding example.

### AlphaPart

AlphaPart is the first free implementation of the method for partitioning breeding values and genetic trends. The method and the package are valuable for deciphering and quantifying the sources of genetic gain in breeding programmes. The package is easy to use, since it streamlines the partitioning analysis into a few lines of R code. AlphaPart presents a holistic tool to perform a partitioning analysis, from preparing the input data - such as manipulating the pedigree data - to handling of results and plotting. The partitioning step is fast, even for large pedigrees, since the main partitioning function is recursive and implemented in C++.

AlphaPart is aimed at the researchers who are interested in quantifying the sources of genetic gain in their breeding programmes either to understand the dynamics of genetic gain, improve efficiency, asses the performance of different breeding actions, optimize investments etc. Such users should take into account the accuracy of the estimated breeding values and their Mendelian sampling terms, which are driven by the biology of the trait and breeding programme structure.

Our future work on AlphaPart will include extending the partitioning method in three areas. The first extension will utilise genomic information to inform which genome regions drive genetic change and what are sources of specific haplotypes or alleles. The second extension will use the partitioning method to analyse changes in genetic variance in addition to the genetic mean. The third extension will simplify handling of uncertainty of path contributions when working with samples from posterior distributions [2, 8].

### Pig breeding example

The pig breeding example showed the investigative power of the partitioning method and the free AlphaPart implementation. Here we discuss the sources of genetic gain in the two tiers of a pig breeding programme.

By partitioning the genetic trend in a simulated pig breeding programme, we disentangled the observation of some multiplier animals having higher breeding values for some traits compared to the nucleus animals. While larger number of recombinations in the multiplier can potentially reveal more variation and occasional outlying animals, we expect lower breeding values in the multiplier due to time-lag between the nucleus and multiplier. The partitioning revealed that the gene flow from the nucleus into the multiplier was the main source of genetic gain in the multiplier, with the nucleus males contributing the most. This was expected due to nucleus-multiplier gene flow and higher intensity of selection in males.

However, the results also showed that selection in the multiplier can contribute genetic gain in addition to the gene flow from the nucleus. The multiplier outperformed the nucleus for trait 1, because with the 10,500 recorded multiplier animals there was substantial amount of information for accurate multiplier selection that generated additional genetic gain. The partitioning of genetic trend for trait 1 showed that when we used only the nucleus males in the multiplier (MaleFlow100), the multiplier generated additional gain from two sources. First, compared to the nucleus, the contribution of the nucleus males increased because they contributed through the gene flow and through the selection of multiplier females. Second, the selection of multiplier females contributed as well. When we used both the nucleus males and the multiplier males in the multiplier (MaleFlow20), the multiplier generated further gain through a combination of the sources. First was the contribution of the selection of multiplier females and males. In contrast, the contribution of nucleus selection decreased due to the reduced gene flow. This decrease was due to a smaller number of progeny per nucleus male compared to the MaleFlow100 scenario.

On the contrary, trait 2 was not measured in the multiplier and had comparable or smaller genetic trend in the multiplier than in the nucleus. For trait 2 the multiplier animals were selected only on estimated parent average, which resulted in low accuracy selection. In the MaleFlow100 scenario this low accuracy selection resulted in a null contribution of multiplier females to the genetic trend for trait 2 and comparable genetic trends between the nucleus and the multiplier. In the MaleFlow20 scenario with a reduced nucleus-multiplier gene flow this low accuracy selection resulted in the reduced genetic gain for trait 2.

## Conclusion

AlphaPart R package is a freely available software for partitioning breeding values and genetic trends. Use of AlphaPart will help breeders to better understand sources of genetic gain and improve their breeding programmes.

## Declarations

### Availability of data and materials

**Project name:** AlphaPart

**Project home page:** https://cran.r-project.org/package=AlphaPart

**Operating system(s):** Windows, MacOS, Linux

**Programming language:** R & C++

**License: GPL-2** | **GPL-3**

**Any restrictions to use by non-academics: -**

